# YeATSAM analysis of the chloroplast genome of walnut reveals several putative un-annotated genes and mis-annotation of the trans-spliced rps12 gene in other organisms

**DOI:** 10.1101/094441

**Authors:** Sandeep Chakraborty

## Abstract

An open reading frame (ORF) is genomic sequence that can be translated into amino acids, and does not contain any stop codon. Previously, YeATSAM analyzed ORFs from the RNA-seq derived transcriptome of walnut, and revealed several genes that were not annotated by widely-used methods. Here, a similar ORF-based method is applied to the chloroplast genome from walnut (Accid:KT963008). This revealed, in addition to the ~84 protein coding genes, ~100 additional putative protein coding genes with homology to RefSeq proteins. Some of these genes have corresponding transcripts in the previously derived transcriptome from twenty different tissues, establishing these as bona fide genes. Other genes have introns, and need to be manually annotated. Importantly, this analysis revealed the mis-annotation of the rps12 gene in several organisms which have used an automated annotation flow. This gene has three exons - exon1 is ~28kbp away from exon2 and exon3 - and is assembled by trans-splicing. Automated annotation tools are more likely to select an ORF closer to exon2 to complete a possible protein, and are unlikely to properly annotate trans-spliced genes. A database of trans-spliced genes would greatly benefit annotations. Thus, the current work continues previous work establishing the proper identification of ORFs as a simple and important step in many applications, and the requirement of validation of annotations.

## Introduction

Common walnut (*Juglans regia* L.) derives its economic importance for its wood and the high nutritional value of the nuts [1]. The walnut genome sequence was recently obtained from the cultivar Chandler [2]. Also, the complete chloroplast sequence (GenBank: KT963008) was annotated with CpGAVAS [3], revealing 86 protein-coding genes [4].

Previously, the YeATS [5] suite was developed to address assembly artifacts in RNA-seq derived transcriptomes [6], and was applied to identify metagenomes [7,8]. The possibility of mis-annotation of genes (transcript from heat shock protein of the fungi *Cladosporium cladosporioides* has been erroneously annotated as a saffron gene) was demonstrated, emphasizing the need to remove contamination from transcriptomes as an important first step [8]. YeATS was extended (YeATSAM) to demonstrate several important omissions by other tools [9]. Finally, it was proposed that the merged transcripts might bias expression counts [10].

Here, the walnut chloroplast sequence is analyzed to identify additional protein coding genes that are not annotated. In addition to finding ~100 putative genes, some of them occuring in the transcriptome obtained earlier [2], a major finding in this study was the identification of the mis-annotation of the rsp12 gene in several organisms. rsp12 is a trans-spliced gene having three exons - exon1 is located ~28kbp from exon2 and exon3 [11,12]. The mis-annotation is a direct result of the complexity faced by *de novo* annotation methods in detecting such genes.

## Methods

The ‘getorf’ program from the EMBOSS suite [13] was used to obtain the ORFs. As described previously, a BLAST database of protein peptides using ~30 organisms from the Ensembl genome [14] was created [9,10]. This is done to reduce computational times. All ORFs > 60 aa were BLAST’ed to this database [15]. Final verification is done on the ‘nr’ database choosing RefSeq proteins. The transcriptome from twenty different tissues is at http://dendrome.ucdavis.edu/ftp/Genome_Data/genome/Reju/transcriptome/Ttinity_Assembly [2, 5, 7]. Multiple sequence alignment was done using MAFFT (v7.123b) [16], and figures generated using the ENDscript 2.0 server [17]. PHYML (v3.0) was used to generate phylogenetic trees from alignments [18]. All protein structures were rendered by PyMOL(TM) Molecular Graphics System, Version 1.7.0.0. (http://www.pymol.org/). Results reported here are obtained using a simple workstation in a few hours.

## Results

The walnut chloroplast sequence analysis revealed ~10000 ORFs (FILE:genome.chloroplast.orf.fa), with 530 ORFs having > 60 aa (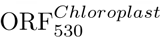, FILE:genome.chloroplast.orf.fa.filterlen.60.fa). There are currently 56036 annotated genes for *J. Regia* in the NCBI database (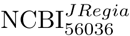, FILE:JRegia.nr.fa). As a corroboration step, 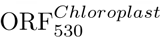 were BLAST’ed [15] to 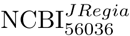, and revealed 84 (and not 86) protein coding genes (FILE:annotated.ncbi.txt). The two missing genes might be due to the 60 aa cutoff applied here.

Next, 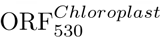 were BLAST’ed to the local BLAST database from ~ 30 organisms (see Methods), and revealed 118 protein coding fragments (FILE:not.annotated.local.ncbi.txt), which have not been annotated. These 118 ORFs were then BLAST’ed to the RefSeq proteins in the ‘nr’ database, resulting in a total of 99 entries (FILE:not.annotated.ncbi.txt). An example is ORF KT963008.1_2812 (125 aa) homologous to the RefSeq protein *(Pharus latifolius,* Accid:YP_008080537.1, Evalue=1E-67) (Fig 1a). This ORF is 100% identical to transcript C43892_G2_I1 identified in previous work [2], and is thus a bona fide gene (Table 3). Some of these ORFs are fragments, and might need manual curation before annotation. For example, three ORFs (KT963008.1_4166, KT963008.1_4182 and KT963008.1_4200) can be merged to create a protein homologous to the RefSeq gene from *Medicago truncatula* (Accid:XP_013455718.1, Evalue=4E-45) (Fig 1b). This gene is ‘’encoded by transcript MTR_4g451295”, and is a bona fide gene at least in *M. truncatula.* Another strategy to exclude pseudo-genes compared the predicted genes to the transcriptome derived from twenty different tissues [2], confirming the presence of at least five real genes (Table 4).

**Figure 1:**
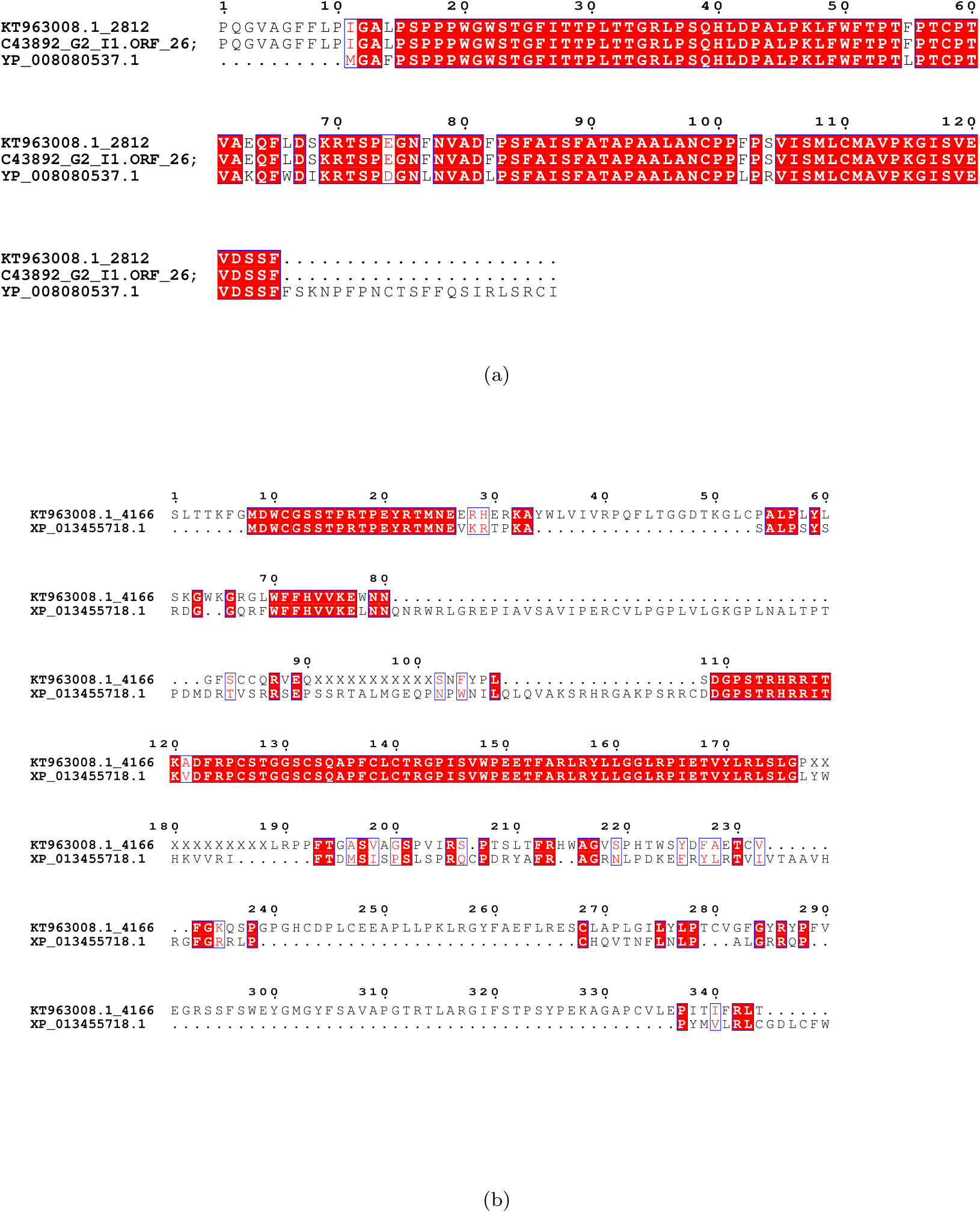
**Chloroplast genes that are not annotated in the current NCBI database:** (**a**) ORF KT963008.1_2812 (125 aa) is homologous to a RefSeq protein (*Pharus latifolius*, Accid:YP_008080537.1, Evalue=1E-67). Furthermore, this ORF is 100% identical to transcript C43892_G2_I1 from a previously derived transcriptome, establishing it as a bona fide gene. (**b**) Manually merging three ORFs to obtain an previously un-annotated gene: The three ORFs are KT963008.1_4166, KT963008.1_4182 and KT963008.1_4200, which have been merged manually by the insertion of a random number of “X”. This is compared to a RefSeq gene from *Medicago truncatula* (Accid:XP_013455718.1), ‘’encoded by transcript MTR_4g451295” (Evalue=4E-45). Thus, this is not a pseudo-gene, at least in *M. truncatula*.

An important finding in the current study is the mis-annotation of the ribosomal rps12 gene in several organisms (Table 1, Fig 2). The rps12 gene has three exons - exon2 and exon3 are proximal, while exon1 (38 aa) is distant, and the complete protein is assembled through trans-splicing [11] (Table 2). A pentatricopeptide repeat protein facilitates the trans-splicing of rps12 [19] (Fig 3). It is not surprising that automated annotation tools will pick up the closest ORF with a start codon prior to exon2 to complete a theoretically possible protein sequence.

**Table 1:** **Enumerating mis-annotated rps12 genes in the NCBI RefSeq database:** The bona fide rps12 gene is 124 aa long, and is identified correctly through conceptual translation. rps12 is a trans-spliced gene with three exons - exon1 is ~28kbp away from exon2 and exon3. Automated methods identify a ORF in the proximity of exon2 with a start codon, leading to a mis-annotated gene. Gnomon is the method for NCBI eukaryotic genome annotation pipeline [20]. JCVI Eukaryotic Genome Annotation Pipeline uses the EVidenceModeler (EVM) [21]. Accid: Accession id.

| Accid | Organism | Length | Method | Date |
| --- | --- | --- | --- | --- |
| ALO71482.1 | *Juglans regia* | 124 | conceptual translation | 20-MAY-2016 |
| YP_007375022.1 | *Quercus rubra* | 124 | conceptual translation | 18-JAN-2013 |
| YP_008963715.1 | *Liquidambar formosana* | 124 | conceptual translation | 20-DEC-2013 |
| XP_015867265.1 | *Ziziphus jujuba* | 143 | Gnomon | 24-MAR-2016 |
| XP_015898012.1 | *Ziziphus jujuba* | 143 | Gnomon | 24-MAR-2016 |
| XP_016724328.1 | *Gossypium hirsutum* | 135 | Gnomon | 18-MAY-2016 |
| XP_012575527.1 | *Cicer arietinum* | 212 | Gnomon | 08-JUN-2015 |
| XP_013443673.1 | *Medicago truncatula* | 159 | EVM, PASA | 25-AUG-2015 |

**Figure 2:**
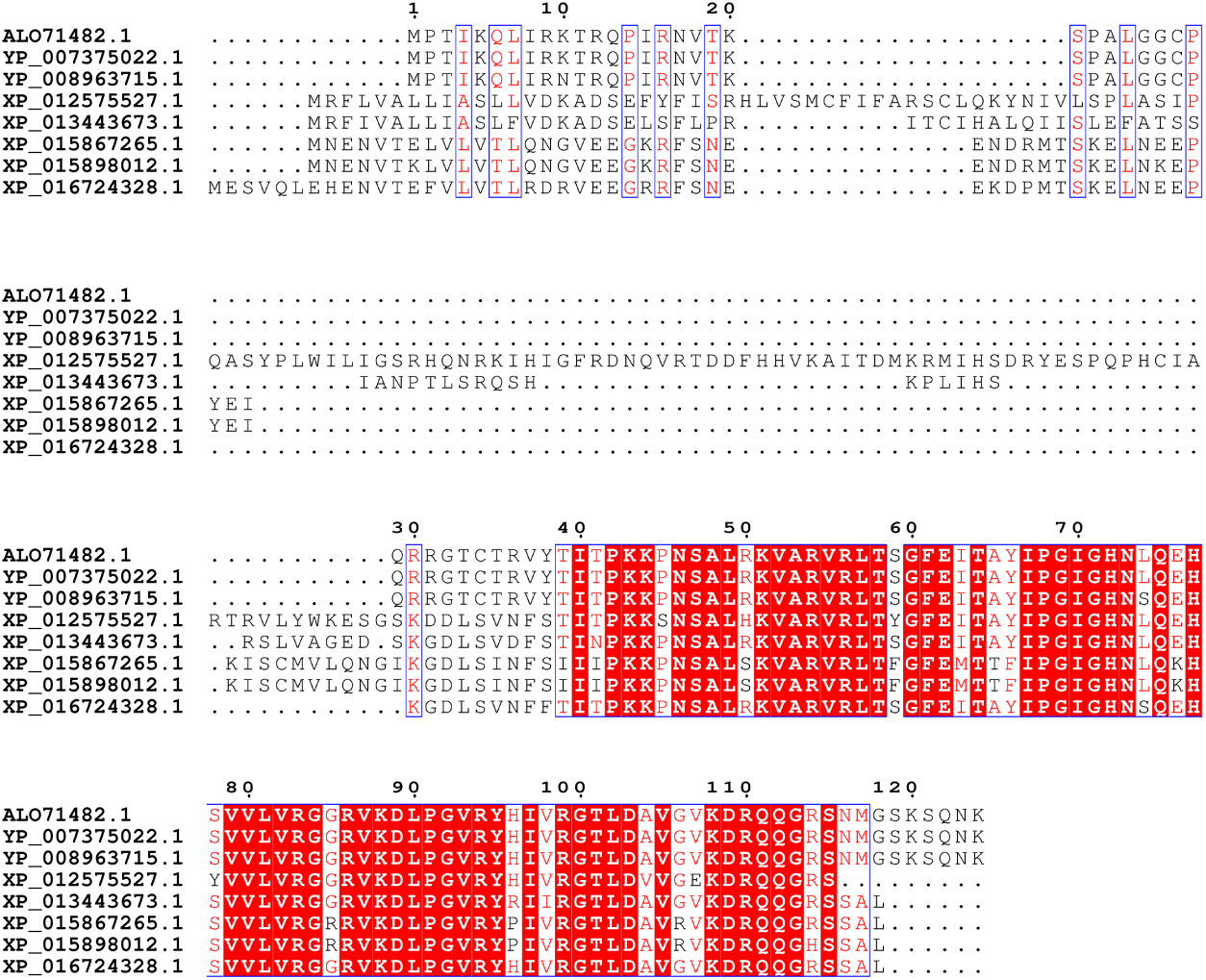
**Multiple sequence alignment of rps12 genes, both bona fide and mis-annotated:** rps12 is a trans-spliced gene with three exons - exon1 is ~28kbp away from exon2 and exon3. ALO71482.1, YP_007375022.1 and YP_008963715.1 are the bona fide rsp12 genes (124 aa long) with exon1 (38 aa long) ending in the sequence “RGTCTRVY”. Mis-annotated genes have chosen ORFs prior to exon2 that lead to a theoretically possible gene.

**Table 2:**
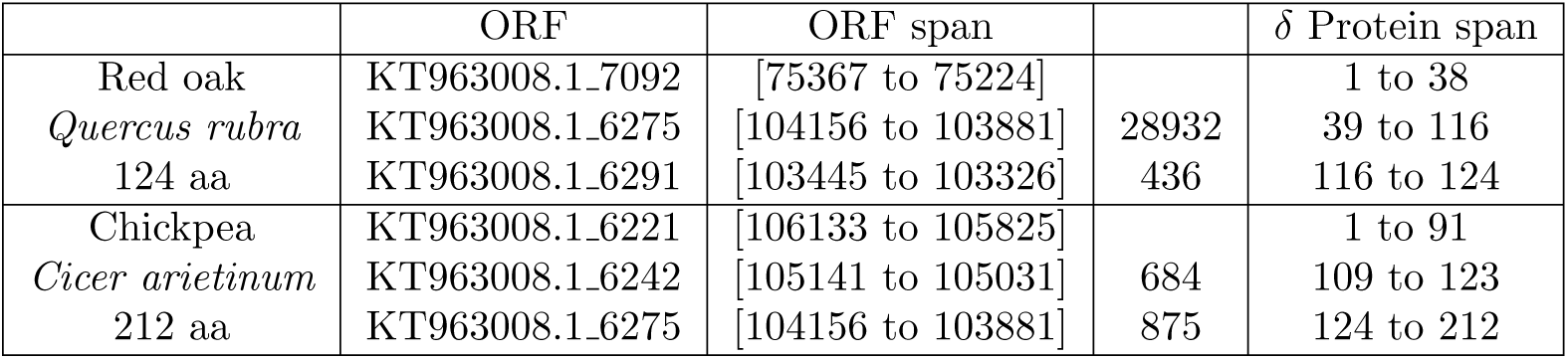
**Mis-annotation of rps12 gene:** The rps12 protein has three exons - exon1 is ~ 28kbp away from exon2 and exon3, and is assembled into the mature protein by trans-splicing [19]. There are other theoretically possible protein sequences that can be assembled by replacing exon1 with a different ORF (KT963008.1_6221 and KT963008.1_6242 in the current case) proximal to exox2.

**Figure 3:**
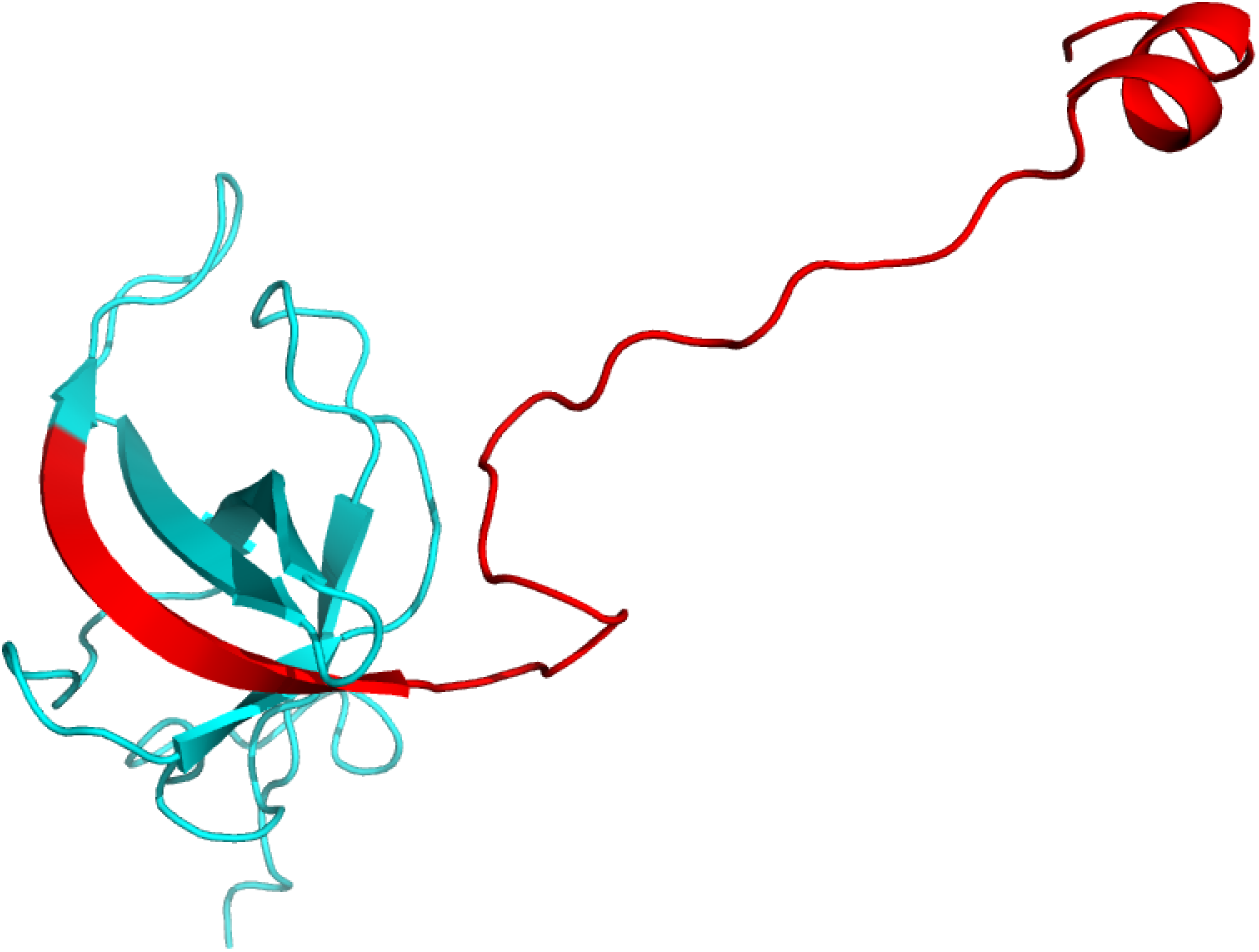
**Structure of rps12 from spinach chloroplast 30s subunit** (**PDBid:3BBN** [12], **chain L:**) rps12 is a trans-spliced gene with three exons - exon1 is ~28kbp away from exon2 and exon3. Exon1 is 38 aa long, and is marked in red.

**Table 3:**
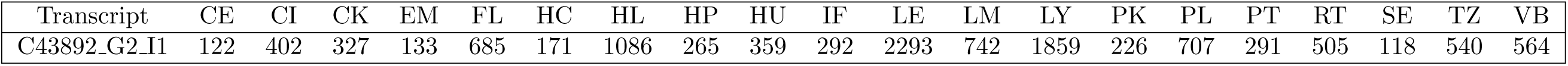
**Expression counts of transcript C43892_G2_I1 which is identical to the ORF KT963008.1_2812**: This gene is not annotated, and is homologous to a RefSeq protein (*Pharus latifolius*, Accid:YP_008080537.1, Evalue=1E-67) These are raw counts, and are not normalized. The abbreviations are provided in FILE:20LIBS.pdf.

**Table 4:** **Comparing predicted ORFs in the walnut chloroplast to the transcriptome**: It can be seen there are at least five bona-fide genes. TRS: transcript id from the RNA-seq transcriptome derived previously [2].

|  | ORF | RefSeq id | Description | TRS |
| --- | --- | --- | --- | --- |
| 1 | KT963008.1_1240 | ERN11400.1 | hypothetical protein *A. trichopoda* | C41199_G1.I1 |
| 2 | KT963008.1_4328 | OIW21822.1 | hypothetical protein *L. angustifolius* | C54203_G1.I1 |
|  | KT963008.1_6215 | OIW21822.1 | hypothetical protein *L. angustifolius* | C54203_G1.I1 |
| 3 | KT963008.1_2812 | YP_008080537.1 | orf137 (chloroplast) *P. latifolius* | C43892_G2.I1 |
|  | KT963008.1_4790 | YP_008080537.1 | orf137 (chloroplast) *P. latifolius* | C43892_G2.I1 |
| 4 | KT963008.1_2840 | NP_817236.1 | ORF58d *P. koraiensis* | C43892_G2.I2 |
|  | KT963008.1_4818 | NP_817236.1 | ORF58d *P. koraiensis* | C43892_G2.I2 |
| 5 | KT963008.1_7519 | DAA50039.1 | TPA: hypothetical protein *Z. mays* | C62693_G1.I1 |

## Discussion

ORF based annotation is trivially simple. Yet, as the current manuscript shows, several protein coding genes remain un-annotated in the walnut chloroplast. Comparison to transcriptomes enables further corroboration of non-pseudo genes. On the other hand, *de novo* identification of trans-spliced genes through automated methods is almost impossible. Mis-annotation of the rps12 gene by automated methods need to be purged before they become mainstream. Also, a database of trans-spliced genes would greatly facilitate their proper annotation. Future work in the YeATS suite will address the automated stitching of spliced genes, identified through fragmented ORFs.

